# CMS4-focused multi-omic integration enhances antigen target identification in colorectal cancer

**DOI:** 10.64898/2026.05.04.722755

**Authors:** Eleonore Fox, Lea Meunier, Solene Weill, Guillaume Appe, Abdelkader Behdenna, Lucas Hensen, Celia Lafond, Akpeli Nordor, Camille Marijon

## Abstract

Colorectal cancer (CRC) remains a major cause of cancer mortality, with limited options for poor-prognosis subtypes such as CMS4. Antigen-targeted therapies show promise but tend to fail due to inadequate target selection and insufficient patient stratification. Effective prioritization requires large harmonized data capturing CRC heterogeneity – a resource that is currently lacking. To address this need, we built a harmonized multi-omic CRC knowledge base and applied a scalable discovery pipeline to identify antigen targets specifically associated with CMS4 biology and with strong translational potential.

We constructed a harmonized CRC atlas by integrating 79 transcriptomics datasets (5,033 tumors, 161 normal samples) using proprietary AI-powered data scouting, integration, and curation technologies. Consensus Molecular Subtypes (CMS) were inferred to capture CMS4-specific expression patterns and this atlas was then combined with 3 bulk RNA-seq reference datasets, 2 single-cell atlases, and 8 protein annotation databases to form a unified multi-omic CRC knowledge base of unmatched scale. From this integrated system, we identified genes differentially expressed in CMS4 patients encoding druggable cell-surface proteins, which we then prioritized using a weighted efficacy- and safety-based scoring model.

We identified 236 CMS4-enriched candidates, including 124 not detectable at the CRC-wide level, demonstrating the added resolution gained through subtype stratification. Recovery of known investigational CRC (LGR5, MET, TACSTD2) and CMS4-associated targets of clinical emerging interest (PDGFRB, ALK5/TGFBR1, FAP) support the biological and methodological validity of our approach.

Benchmarking against thresholds from FDA-approved pan-cancer targets and terminated trials identified 32 candidates with comparable or superior therapeutic profiles. Among these, 11 were enriched for CMS4-defining pathways, including epithelial–mesenchymal transition, angiogenesis, and stromal invasion, and 5 showed strong profile similarity to established CRC and CMS4 benchmarks. After extensive data exploration, particularly promising candidates were shortlisted for further validation.

This work shows that CMS4-focused molecular stratification, when combined with an unprecedentedly large harmonized multi-omic knowledge base, yields a refined set of antigen candidates with enhanced specificity, safety, and biological relevance. The prioritized targets illustrate the power of subtype-resolved discovery to uncover clinically actionable insights. Our pipeline’s modular design can extend to other tumor contexts, offering a robust foundation for accelerating targeted therapy development.

## INTRODUCTION

Colorectal cancer (CRC) is the third most commonly diagnosed malignancy and the second leading cause of cancer-related mortality worldwide (1), accounting for 6.1% of new cancer cases and 9.2% of cancer deaths (2). Despite advances in screening and therapeutic modalities, advanced disease remains associated with poor outcomes, with 5-year overall survival below 15% in metastatic CRC (3). CRC’s molecular heterogeneity limits durable therapeutic benefit and underscores the urgent need for biologically informed precision strategies to identify and effectively target actionable vulnerabilities (4).

The Consensus Molecular Subtypes (CMS) define four biologically and clinically distinct CRC subgroups. Among these, CMS4 tumors are characterized by mesenchymal transition, stromal remodeling, angiogenesis, TGF-β pathway activation (5) and exhibit poor prognosis alongside limited response to conventional therapies (Figure S2C-D). Although CMS classification has improved understanding of CRC biology, it is not routinely used to guide therapeutic decisions. Nonetheless, its subtype-specific molecular features offer an opportunity to identify actionable targets in high-risk populations such as CMS4 and support the integration of molecular patient stratification in clinical practice (6).

While antigen-targeted therapies have transformed oncology, their impact remains limited by antigen heterogeneity, insufficient tumor specificity and on-target, off-tumor toxicity (7). The identification and selection of novel antigen targets highly expressed in tumor cells, minimally expressed in essential healthy tissues, and structurally accessible to therapeutic intervention remains a major challenge (8,9).

Antigen target discovery methods found in the literature historically relied on time-consuming experimental screens (10), single-cohort analyses, and limited integration of omic, clinical, or drug development evidence (11–13). Candidate lists are consequently often extensive, weakly prioritized, and insufficiently informed by safety profiles or real-world therapeutic benchmarks (14). This inefficiency drives high attrition in oncology drug development, where nearly 90% of clinical programs fail, most frequently due to inadequate efficacy or unacceptable toxicity (15). While public repositories provide access to large and diverse omic datasets, their potential remains underexploited owing to poor harmonization and inconsistent clinical annotation (16). Data-driven strategies integrating multi-omic, clinical, and translational evidence allow for improving early-stage target selection and mitigating downstream development risk (17). Large-scale harmonization and aggregation of public datasets increases patient representation and diversity, statistical power, and enables analysis of tumor biology by molecular subtype (18).

Here, we present a multi-omic framework for antigen target discovery in colorectal cancer, focusing on the poor-prognosis CMS4 subtype. We constructed a large-scale, harmonized CRC knowledge base integrating transcriptomic, proteomic, genomic, structural, and clinical trial evidence across 16,237 protein-coding genes. Within this infrastructure, we implemented a subtype-informed discovery strategy coupled to a weighted scoring system evaluating predicted efficacy, safety, and therapeutic novelty of potential targets. Scoring thresholds were benchmark-calibrated using FDA-approved antigen-directed therapies alongside discontinued clinical trials to provide translational grounding.

Utilizing this approach, we identified and prioritized biologically and clinically coherent antigen candidates enriched in CMS4 tumors. These results support the feasibility of systematic data integration and benchmark-calibrated prioritization for subtype-specific antigen target discovery in oncology.

## METHODS

### Data acquisition, preprocessing, and harmonization

#### Clinical data curation

All public clinical datasets included in this study were selected and harmonized through a proprietary human-in-the-loop framework combining pattern-based and LLM-based AI models with systematic expert review (unpublished data). Harmonization was achieved by mapping heterogeneously described clinical data onto internally developed, expert-curated ontologies — one per clinical parameter — iteratively refined to reflect evolving biological knowledge and study-specific requirements. Where applicable, these ontologies were aligned with established international standards, including the World Health Organization’s International Classification of Diseases for Oncology (ICD-O) (19) and the Cell Ontology (20); concepts lacking standardized equivalents were defined by expert consensus.

This hybrid curation framework was designed to balance cross-institutional interoperability with the analytical granularity required to support drug discovery and translational research.

#### Microarray data processing

Publicly available colorectal cancer microarray datasets were retrieved from the Gene Expression Omnibus (GEO)(21) (Table S1). Datasets were restricted to the Affymetrix GPL570 platform (HG-U133 Plus 2.0) to ensure technical comparability. Inclusion criteria required biopsy site annotation corresponding to colon, rectum, or rectosigmoid junction, and sample type annotated as primary tumor or normal tissue from healthy donors. Raw CEL files were processed in R using the OligoR Bioconductor package(22). Robust Multi-array Average (RMA) normalization was applied for background correction, quantile normalization, and log2 transformation.

Preprocessed datasets were merged into a single atlas. Batch effects were corrected using ComBat (pyComBat implementation), with sample type included as a biological covariate to preserve tumor-normal differences while removing technical variability (23). This correction strategy was enabled by the presence of datasets containing both healthy and tumor samples, which anchored the biological covariate signal across batches.

#### Bulk RNA sequencing data processing

Bulk RNA sequencing count data were obtained from multiple sources: tumor samples from The Cancer Genome Atlas (TCGA Data Release 36.0) via the Genomic Data Commons ([GDC](https://portal.gdc.cancer.gov/)) portal (24), normal tissues samples from the Genotype-Tissue Expression ([GTEx v8](https://www.gtexportal.org/home/downloads/adult-gtex/bulk_tissue_expression)) project (25), and colorectal cancer cell lines from the Cancer Cell Line Encyclopedia ([CCLE 2019](https://depmap.org/portal/data_page/?tab=allData)) (26). Cell lineage annotations for CCLE samples were harmonized to ensure concordance with colorectal tissue classification. When available, cell line molecular subtype classification was assigned based on Cellosaurus annotations (27). Raw count matrices were normalized using the trimmed mean of M-values (TMM) method to ensure comparability across cohorts.

#### Somatic mutation data

Somatic mutation data were retrieved from TCGA (Data Release 36.0) Mutation Annotation Format (MAF) files using TCGAbiolinks (28). Only coding non-synonymous variants were selected as they likely introduce amino acid changes that directly impact protein structure and function, making them functionally relevant candidates. Mutation frequencies were computed per indication as the proportion of patients harboring at least one qualifying variant in the gene of interest.

#### Single-cell RNA sequencing data processing

Single-cell RNA sequencing datasets were retrieved from the Gene Expression Omnibus (GEO) for colorectal tumor samples and from Tabula Sapiens for healthy tissues across multiple organs (29). Data processing followed established single-cell best practices in the literature (30) and was implemented using Scanpy (v1.9) in Python (31).

Quality control was performed in three steps. First, ambient RNA contamination was removed using SoupX (32). Second, cells were filtered if they deviated by more than 5 median absolute deviations (MADs) from the sample distribution across standard quality metrics, including percentage of mitochondrial, ribosomal and hemoglobin genes. Doublets were identified and excluded using DoubletFinder (33).

Gene expression counts were normalized using Scran (34), which estimates cell-specific size factors pooled across locally similar cells to account for technical variation while preserving biological signals. Normalized data were log-transformed prior to dimensionality reduction.

Unsupervised clustering was performed using the Leiden algorithm at resolution 0.5, followed by marker gene identification to characterize cluster identities. Cell type annotation was performed by integrating two complementary approaches: CancerFinder(35) to distinguish tumor from microenvironment cells and scTAB(36) for precise cell type prediction. Final labels were harmonized to an internal cell type ontology to enable cross-dataset comparison.

#### Proteomic data processing

Subcellular localization annotations were obtained from the Human Protein Atlas (HPA) (37) and the COMPARTMENTS database (38), and harmonized using Gene Ontology (GO) mapping (39). Plasma membrane localization and associated confidence scores were retained and rescaled to a 0–10 scale. Transmembrane domain, GPI-anchor, and secretion annotations were obtained from UniProt (40).

Protein expression levels in normal tissues were compiled from HPA (37), the Human Proteome Map (41), and ProteomicsDB (42) which provides antibody- and mass spectrometry–based quantification. Tissue names were harmonized to an internal consensus ontology. Expression values were discretized into four categories (0: not detected; 1: low; 2: medium; 3: high) by fitting distributional thresholds approximating a normal distribution. For each gene–tissue pair, the highest reported category across databases was retained to ensure conservative safety modeling. Tissues were classified as vital or non-vital based on predefined clinical criteria.

Protein expression in tumor pathology samples (immunohistochemistry) was obtained from HPA and converted from qualitative categories into a standardized 0–3 scale.

Data were accessed via their respective download portals and API. (HPA: https://www.proteinatlas.org/about/download, Compartments: https://compartments.jensenlab.org/Downloads, ProteomicsDB: https://www.proteomicsdb.org/api)

#### Gene name harmonization

All data sources described above were harmonized at the gene level prior to integration. All gene and protein names were mapped to GENCODE v36 nomenclature to ensure consistency across datasets and external databases. To support this process, a mapping database was constructed from seven reference resources, including GENCODE, Ensembl, UCSC, HGNC, NCBI, HPA, and UniProt, incorporating both current and legacy releases. It integrates stable gene IDs, approved symbols, aliases, and full names.

#### CMS prediction

CMS subtypes can be systematically inferred from transcriptomic data, making them reproducible and information-rich molecular annotations.

For all transcriptomic tumor samples, Consensus Molecular Subtypes (CMS1–CMS4) were assigned using a single-sample predictor based on a proprietary Python reimplementation of the CMSclassifier R package (28). Classification was based on a nearest-centroid approach, in which each sample was correlated to predefined subtype centroids and assigned to the subtype with the highest similarity score. Samples not meeting predefined confidence thresholds were excluded from subtype-specific analyses.

Predicted CMS labels were integrated into the harmonized metadata and used for downstream subtype-specific analyses.

#### Pathway annotation

Hallmark, GO, Biocarta and KEGG pathway annotations were obtained from the Molecular Signatures Database (MSigDB v2023.1.Hs(29)). Immune and stromal signatures were obtained from ESTIMATE, a method to infer the fraction of stromal and immune cells in tumour samples (44).

### Target identification pipeline

#### Differential gene expression analysis

Differential expression analyses were performed on the microarray atlas using a Python implementation of limma (30). Two comparisons were conducted: (i) all primary colorectal cancer (CRC) tumors versus normal colorectal tissues and (ii) CMS4 tumors versus normal colorectal tissues. Genes presenting a Benjamini-Hochberg adjusted P value below 0.05 and a log2 fold change above 1 were considered significantly overexpressed.

#### Proteomic expression quality control in normal tissues

To enable safety modeling, genes lacking protein-level expression evidence in healthy tissues were excluded. Genes were retained if protein expression was reported in at least one reference database, ensuring availability of data for downstream safety assessment.

#### Cell surface accessibility filtering

To prioritize targets compatible with antibody- and cell-based therapies, candidates were restricted to plasma membrane–localized proteins. Proteins annotated as plasma membrane–localized in at least one curated source were retained.

Structural filtering was subsequently applied to ensure extracellular accessibility. Only proteins containing at least one predicted transmembrane domain or a glycosylphosphatidylinositol (GPI) anchor were retained.

### Target prioritization framework

#### Scoring dimensions

Targets were prioritized using a quantitative scoring framework integrating three dimensions: efficacy, safety, and novelty.

- Efficacy captures tumor specificity, biological relevance, and translational potential within the indication and, where applicable, the CMS4 subtype. This dimension incorporates tumor expression intensity and prevalence, association with patient outcomes, mutation frequency, hallmark of cancer pathway involvement, and expression in disease-relevant cellular models.
- Safety assesses on-target/off-tumor toxicity by analyzing expression across healthy tissues and cell populations at both transcriptomic and proteomic levels. This includes assessment of expression in vital organs, distribution across normal cell types, and relative tumor-to-normal differential expression.
- Novelty reflects prior exploration in oncology and targeted therapy development, based on structured bibliometric analyses. This dimension provides contextual insight into the competitive landscape and translational maturity.

#### Score normalization and aggregation

For each target, individual parameters’ values were normalized and converted into standardized scores ranging from 0 to 10 using empirical thresholds derived from the distribution across all evaluated candidates. Dimension-specific scores (efficacy, safety, novelty) were calculated as weighted means of their respective parameters, and a global prioritization score was computed as a weighted composite of the three dimensions. Scores do not have units and therefore allow direct comparison of the different values between targets, with scores <5 considered low. They can also be ordered in various ways depending on the interests and therapeutic modality, thereby prioritizing the exploration of the most promising targets.

Weights were either user-defined or set to default values. Default weights were established through benchmarking against FDA-approved antigen-targeted cancer therapies, established therapeutic targets, and programs discontinued due to safety or efficacy limitations.

#### Efficacy, safety, and novelty score computing

Efficacy and safety scores were computed using a structured parameterization framework detailed in the associated <u>Excel file</u> (Supplementary Data). The file specifies all variables, normalization procedures, weighting schemes, and aggregation rules applied for dimension-specific and global score calculation.

### Identification of benchmark therapeutic targets

#### Sources

Previously explored therapeutic targets were identified through structured interrogation of regulatory and pharmacological databases. ChEMBL was used as the primary source to retrieve drug names and associated pharmacological information, including target identity and mechanism of action. Extracted drug and target annotations were subsequently used to query ClinicalTrials.gov for indication-specific clinical data. Patsnap was interrogated to complete missing information and cross-validate entries, leveraging its standardized data structure to improve consistency across sources. Drug approval status was confirmed using the FDA Drug Approvals and Databases resource.

#### FDA-approved antigen-directed therapies

FDA-approved antigen-directed therapies were identified through structured queries of regulatory and pharmacological databases. Eligible agents were required to (i) be targeted therapies, (ii) act against cell-surface protein antigens overexpressed in malignant cells, (iii) be approved for at least one oncologic indication, and (iv) belong to ATC classes L01F, L01XL, L01XX, or V10X.

A total of 9 target and 18 target–indication pairs met these criteria (Sup Table 4).

#### Well-established therapeutic targets

Well-established therapeutic targets were defined as targets that have been evaluated in clinical trials. A search was conducted on ClinicalTrials.gov using the condition “*Indication*”, the respective indication name, and the interventions “CAR” or “ADC”. The resulting studies were manually screened to identify therapeutic targets relevant to each indication. Trials of all phases involving chimeric antigen receptor T cells (CAR-T) or antibody–drug conjugates (ADCs) were included, while basket trials were excluded. Only trials that were indication-specific or explicitly mentioned the indication of interest in the trial title were considered.

#### Programs discontinued due to safety or efficacy

Clinical trials discontinued due to safety or efficacy issues were identified by manually reviewing studies for the relevant indications on ClinicalTrials.gov with the status “Suspended” or “Terminated”. Trials were classified as discontinued due to safety and/or efficacy based on the reported termination reason.

To facilitate scalable screening, keyword-based filtering was applied to the termination reasons. Safety-related keywords included *safety, toxicity, toxicities*, and *DSMB/DSMC*. Efficacy-related keywords included *efficacy, DSMB/DSMC, exposure, statistical success, ineffective, tumor response*, and *nonsatisfactory clinical benefit*.

To reduce false positives, trials containing termination reasons related to operational or strategic factors (e.g., *sponsor decision, funding, slow enrollment, poor recruitment, drug supply, protocol changes*, or *corporate strategy*) were excluded.

#### Benchmark calibration

Efficacy and safety score thresholds were calibrated using the identified panel of FDA-approved antigen-targeted oncology therapies. When cancer type-specific data were unavailable, score calculations were adjusted to prevent penalization due to missing information.

Across the benchmark set, the mean efficacy score was 5.45 (median 5.8) and the mean safety score was 5.1 (median 4.85). An efficacy and safety threshold ≥5.0 was therefore defined as reflective of profiles comparable to clinically validated antigen-directed therapies, with slightly lower cutoffs considered permissive alternatives. Candidates exceeding defined thresholds were classified as meeting benchmark-derived translational criteria.

#### Quantification and statistical analysis

All analyses were performed using Python (v3.11) and R (v4.2).

## RESULTS

### Construction of a clinically oriented multi-omic CRC cancer knowledge map

To identify antigen targets in CRC, we constructed an integrated knowledge base using multiple resources that provide complementary data on colorectal cancer biology. This database enables the evaluation of candidate proteins against key criteria for targeted therapy, including tumor specificity, membrane accessibility, expression in healthy tissues, and clinical evidence (Figure 1A).

**Figure 1:**
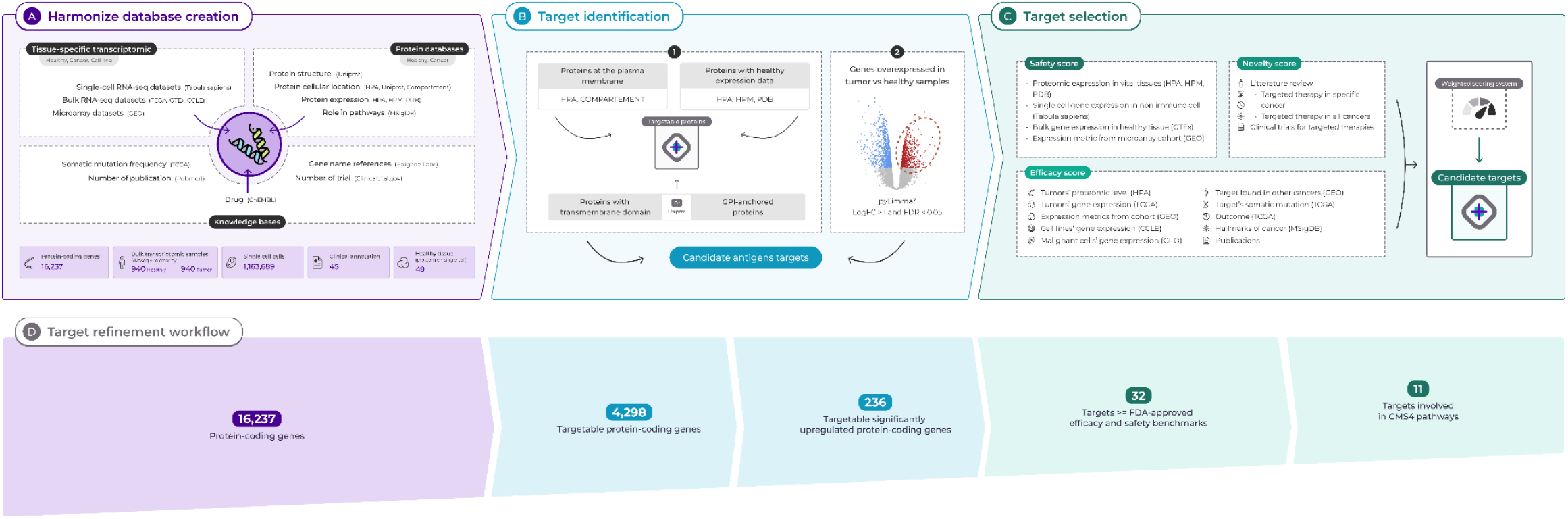
Multi-omic colorectal cancer target discovery workflow: (A) Knowledge base construction, integration of diverse data sources into a unified database; (B) Target identification based on predefined criteria; (C) Multi-dimensional ranking of candidates and selection of top targets; (D) Funnel chart illustrating the progressive refinement and prioritization of targets across workflow stages.

To assess target expression at the population level, we constructed a large-scale transcriptomic atlas comprising 5,194 tumour and normal colorectal samples from 77 microarray studies (GEO) using our proprietary AI framework (Table S2). Expression data were harmonized into a unified format, and associated clinical metadata were curated at scale through an AI-assisted, human-in-the-loop framework combining automated extraction with expert review (Figure S1) (unpublished data). The GEO cohort included 5,033 colorectal cancer patients (mean age 65.0 ± 12.8 years; 56.4% male among those with available data), with tumours spanning stages I–IV and arising predominantly from the colorectum or colon, though sex, age, and stage annotations were missing for approximately half of the patients. This atlas complements RNAseq expression profiles from the TCGA colorectal adenocarcinoma cohort (COAD/READ), which comprised 637 patients with near-complete clinical annotation (mean age 66.4 ± 12.7 years; 53.4% male), tumours predominantly at stage II (38.0%), and a colon-predominant biopsy distribution (73.8%) (Table S3). While both cohorts are broadly comparable in age and sex distribution, the TCGA cohort is skewed towards earlier-stage disease relative to the GEO cohort. Together, the GEO atlas increases the number of available samples by approximately 650% relative to TCGA alone, substantially expanding patient representation and molecular heterogeneity. Target expression was then examined across commonly used preclinical models using transcriptomic profiles from colorectal cancer cell lines (CCLE), enabling comparison between experimental models and patient tumors.

Gene expression was further contextualised within the cellular landscape of healthy tissues using single-cell transcriptomic data, providing a reference for baseline expression across physiological cell types (Tabula Sapiens). Comparative expression across cell types within tumours and the tumour microenvironment was assessed using single-cell RNA sequencing datasets from colorectal cancer samples (GEO), allowing expression profiling in malignant and non-malignant compartments. The genomic landscape of candidate targets was characterized by examining somatic mutations and copy-number alterations in colorectal cancer (TCGA). Protein-level characterization relied on the integration of proteomic and structural resources documenting tissue expression, protein structure, function, and subcellular localisation (Human Protein Atlas, Human Proteome Map, Protein Data Bank, UniProt, and COMPARTMENTS), while curated gene-set collections provided pathway-level functional annotation (MSigDB). Finally, the therapeutic landscape surrounding candidate targets was surveyed using resources capturing drug–target associations, clinical development status, and related literature, although these datasets remain distributed across heterogeneous formats that limit direct interoperability (ChEMBL, ClinicalTrials.gov, PubMed).

All the resources included are harmonized into a unified, structured database encompassing 16,237 protein-coding genes, each annotated with 10 standardized features across 4 harmonized data layers. It enables direct comparison under uniform antigen-specific constraints.

Building upon this integrative foundation, we develop a subtype-informed evaluation framework that enables the consistent exploration of targets from clinically validated ones to biologically supported but underexplored targets. Applying this framework, we prioritized antigen targets specifically enriched in the high-risk mesenchymal CMS4 subgroup of colorectal cancer (Figure 1C).

### CMS4-informed identification of cell-surface antigen targets

To identify candidate targets, we first determined the targetable proteins (Figure 1B). We began with plasma membrane proteins and proteins showing structural features consistent with extracellular exposure, such as a transmembrane domain or GPI anchoring. To address the risk of on-target/off-tumour toxicity, we further selected targets for which proteomic expression data was available in healthy tissues. This left us with a pool of 4,298 targetable proteins (Figure 2A).

**Figure 2:**
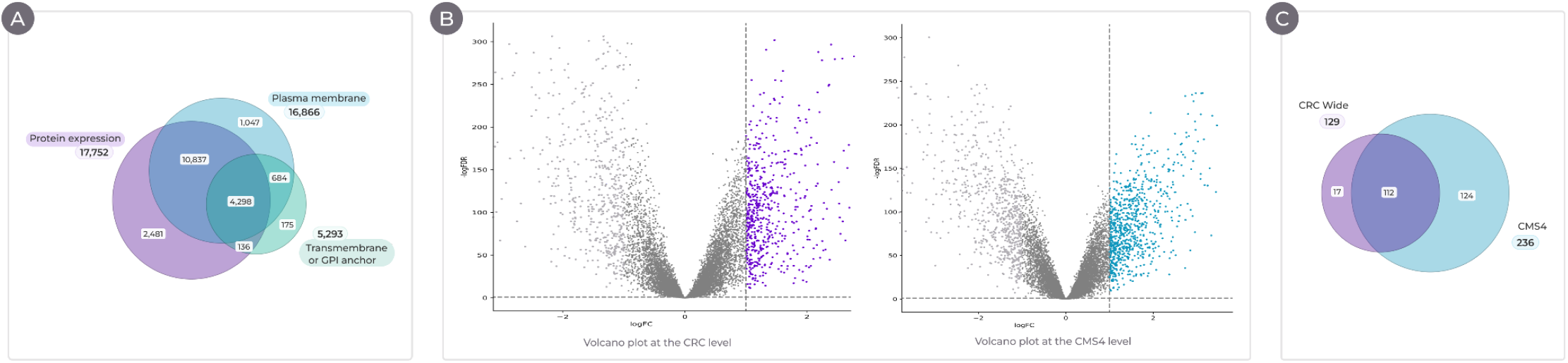
(A) Venn diagram illustrating the targetable surface protein space. (B) Volcano plots showing differential expression results at the CRC level (left) and the CMS4 level (right). (C) Venn diagram depicting the overlap of potential targets identified in the CRC-level and CMS4-level analyses.

Candidates have to be subsequently restricted to the ones overexpressed in the population of interest, CRC and CMS4. To identify CMS4-focused targets, we first inferred molecular subtypes from transcriptomic data using a validated classifier to ensure robust subtype assignment. CMS4 tumors represented 12% of the atlas, consistent with the distribution reported in the seminal study (Figure S2A) (5). Then, we performed a differential expression analysis comparing CMS4 primary tumors to normal colorectal tissues from healthy donors, and identified 836 significantly upregulated genes in this subpopulation (FDR < 0.05, log2 fold change ≥ 1), and 580 upregulated genes by comparing all CRC primary tumors to normal colorectal tissues (Figure 2B). By integrating overexpressed genes with targetable proteins, we uncovered 129 potential CRC targets and 236 for CMS4.

Interestingly, 124 of the 236 CMS4 targets were not identified at the colorectal cancer level (Figure 2C), indicating that subtype-restricted analysis reveals targets otherwise overlooked. The workflow also recovered established and investigational CRC antigens both at the CMS4-specific and at CRC-wide level, including LGR5, MET, and TACSTD2 (TROP2) (46–48), supporting the robustness and translational validity of the discovery pipeline (see complete list of CMS4 well-established targets in Table S5). Moreover, the identification of subtype-coherent targets such as PDGFRB, TGFBR1 (ALK5), and FAP aligns with the mesenchymal, stromal-rich, and TGFβ-driven biology that characterizes CMS4 tumors, demonstrating concordance between molecular subtype biology and candidate antigen selection (Figure S2B) (49,50).

### Multi-dimensional prioritization and benchmark-calibrated target selection

The 236 CMS4-enriched cell-surface antigen candidates were subsequently prioritized. Each candidate was scored across three clinically relevant dimensions: efficacy, safety, and therapeutic novelty (Figure 3A). Scoring incorporated both CRC-wide metrics and CMS4-specific features to ensure that the prioritization captured general indication relevance as well as subtype-specific biology. Detailed scoring methodology is described in the Methods under “Scoring dimensions”.

**Figure 3:**
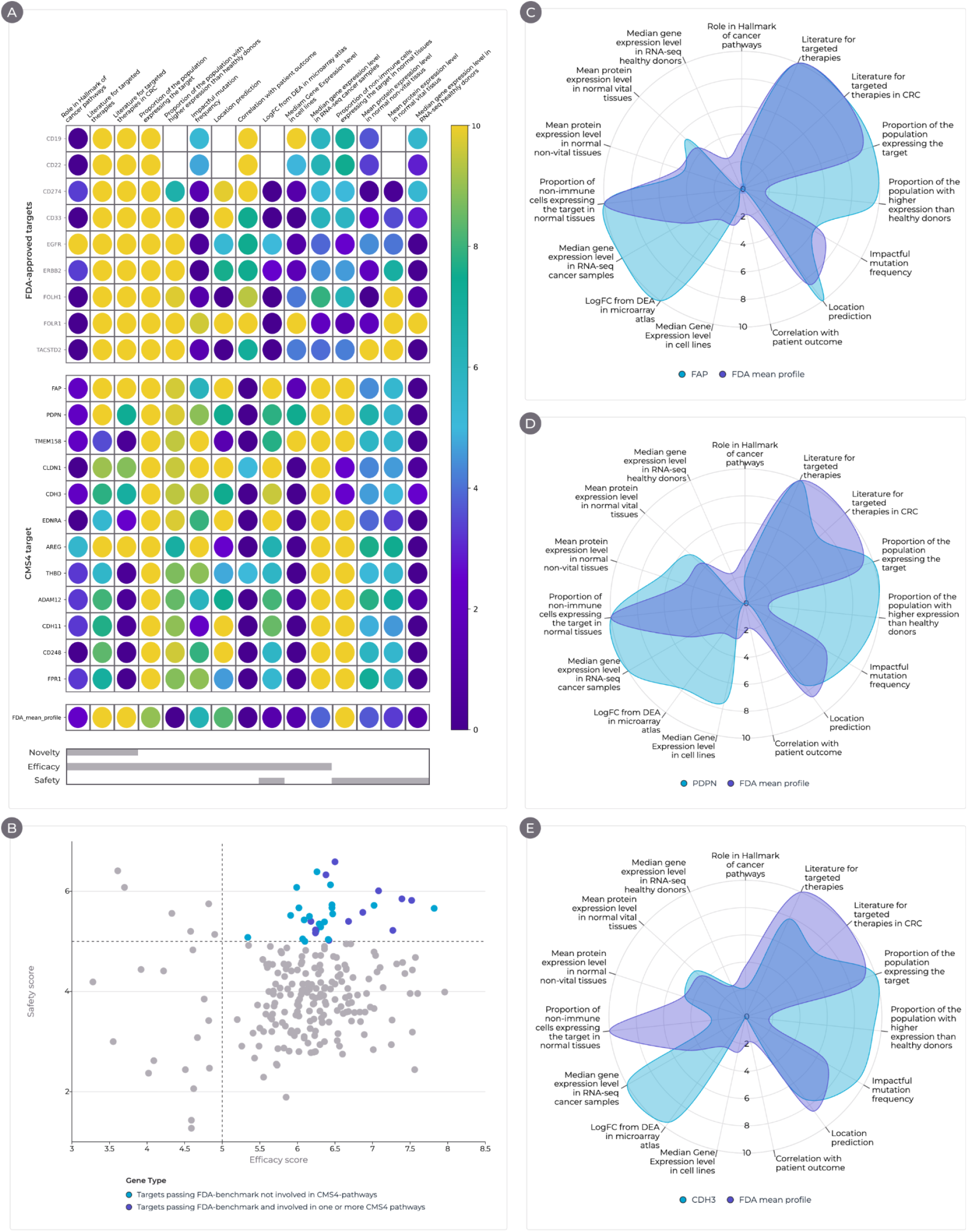
(A) Heatmap illustrating target prioritization scores for FDA-approved targets, CMS4 targets passing benchmarks and involved in CMS4 pathways, and the mean FDA profile. (B) Scatter plot showing target efficacy *versus* safety scores; targets in the upper left quadrant represent the safest and most effective candidates passing FDA benchmarks. (C) Radar chart comparing FAP profile (D) PDNP profile and (E) CDH3 profile against the FDA-mean profile.

To contextualize targets’ potential, efficacy and safety thresholds were calibrated against benchmark profiles derived from FDA-approved antigen-targeting therapies and from clinical programs discontinued due to safety or efficacy limitations. FDA-approved targets showed high transcriptomic tumor expression (with a median of 111 Trimmed mean value of M (TMM)) and broad tumor prevalence (98% of samples), with heterogeneous (but generally limited) protein expression across healthy tissues, consistent with an acceptable safety profile (Figure 3A). This calibration enabled interpretation of candidate scores relative to clinically validated and unsuccessful targets, providing an anchored reference for ranking.

Amongst the antigen candidates, 32 candidates displayed profiles comparable to or exceeding benchmarks derived from approved antigen-directed therapies (Figure 3B). These targets showed high tumor expression in TCGA CMS4 samples (median ≥124 TMM), and were detected in 99% of tumors. Protein-level expression in healthy tissues remained low, consistent with a manageable risk of on-target, off-tumor toxicity. Half of the candidate targets were associated with at least one hallmark of cancer pathway, and 11 were enriched in CMS4-related pathways, including epithelial–mesenchymal transition, angiogenesis, and stromal remodeling, consistent with the biology of this subtype (Table S6).

Among the prioritized candidates, several have been explored in targeted therapy programs outside colorectal cancer (CRC), supporting their translational potential while underscoring their relative novelty in this context. For example, TGFBR1 has been targeted by the TGFβ type I receptor kinase inhibitor vactosertib (in combination with pomalidomide) in a phase 1b trial in relapsed/refractory multiple myeloma (NCT03143985) (51), and MMP14 by BT1718, a first-in-class bicycle toxin conjugate, in a phase I/II dose-escalation study in patients with advanced solid tumours (NCT03486730) (52). The remaining candidates are supported by prior CRC literature, reflecting a balance between innovation and existing validation (ref). Three were ultimately prioritized for further investigation: Fibroblast Activation Protein Alpha (FAP), Podoplanin (PDPN) and Cadherin 3 (CDH3).

Within CMS4 patients, FAP demonstrated elevated efficacy and safety scores, exceeding the median by one standard deviation in both dimensions (Figure 3C), underscoring its therapeutic potential. This surface serine protease is highly overexpressed in cancer-associated fibroblasts (CAFs) across >90% of epithelial tumors, including colorectal cancer (CRC), while being largely absent in normal adult tissues (53–55). Its status as an actionable target was further validated by external databases such as Patsnap and Pharos, which identified active ligands and over 100 clinical trials, including 8 in CRC (56).

PDPN, a mucin-like transmembrane glycoprotein widely upregulated in CAFs and tumor cells across aggressive cancers (57,58) was identified at both the indication and sub-indication levels (Figure 3D). It achieved the highest composite score, surpassing the benchmark defined by FDA-approved antigen-targeting therapies in CMS4 CRC. PDPN-positive myofibroblasts are a hallmark of TGF-β-driven stroma (59) and have been associated with immunotherapy resistance (60).

CDH3 was only identified at the CMS4 level, setting it apart from pan-CRC targets (Figure 3E). Despite an early-phase dose escalation trial’s premature termination due to insufficient anti-tumor activity (61), CDH3 has since emerged as a serum biomarker in metastatic colorectal cancer (62) and as a candidate target for combination immunotherapy (63,64), underscoring its ongoing translational relevance.

Together, this benchmark-calibrated, multi-dimensional prioritization strategy defines a focused, biologically coherent, and clinically actionable set of antigen targets tailored to the high-unmet-need CMS4 colorectal cancer population.

## DISCUSSION

Multiple existing resources provide complementary data on colorectal cancer biology, but differences in structure and annotation prevent their direct integration. They are typically analyzed in isolation, hindering systematic comparison across protein-coding genes. In this work, we addressed this limitation by harmonizing them into a unified, structured database integrated within a unified evaluation framework that enables the consistent exploration of targets from clinically validated ones to biologically supported but underexplored targets.

Subtype-specific analysis reveals targets masked in CRC-wide analyses. This finding underscores the limitations of analyzing CRC tumors as a uniform population, where biological heterogeneity can dilute context-specific signals. CMS4 tumors’ characteristics, such as mesenchymal transition, stromal remodeling, and TGFβ pathway activation, reshape antigen expression landscapes and highlight distinct therapeutic vulnerabilities.

Beyond predefined molecular subtypes, the framework can accommodate alternative stratification strategies based on biologically defined tumor groups or dominant oncogenic processes. This flexibility enables target prioritization across diverse tumor contexts, from discrete molecular subtypes such as CMS4 to tumors defined by heightened activity of specific pathways (TNFα signaling, epithelial–mesenchymal transition, etc.) or specific biomarkers.

A key objective of this work was to move beyond descriptive ranking toward clinically interpretable prioritization. Calibration of efficacy and safety scores against FDA-approved antigen-directed therapies and discontinued programs provided a translational reference framework. Rather than relying solely on internal scoring distributions, this strategy positioned candidate targets relative to real-world therapeutic performance. Survival association informed the predicted safety profile, while tumor specificity and healthy tissue expression helped control on-target off-tumor toxicity risk. Pathway relevance linked candidates to actionable drug mechanisms, and prior clinical exposure provided insight into novelty and existing efficacy or safety signals, thereby enabling multi-dimensional evaluation under drug-development–relevant constraints.

Multiple limitations were identified in this study, including the interpretation of discontinued programs, which remained limited by both the quantity and heterogeneity of publicly available clinical trial data, as reasons for trial termination are often incompletely reported or non-standardized. While efforts have been made to harmonize these data, deeper curation and incorporation of additional patient-level information would improve benchmark calibration. For instance, physical accessibility of tumor cells is rarely documented, yet in immune-excluded or stromal-dense tumors, limited therapeutic penetration may reduce efficacy independently of target expression. In addition, modality-specific benchmarking could also refine scoring, as different therapeutic platforms have distinct biological and structural constraints and not all targets are equally suitable across modalities. However, the limited availability of spatially resolved transcriptomic data and variability in clinical trial reporting currently restrict the implementation of these refinements.

Beyond CMS4-specific findings, the harmonized CRC knowledge base provides an extensible infrastructure for multi-omic antigen target evaluation. By integrating transcriptomic, proteomic, functional, and clinical evidence within a unified scoring framework, it enables rapid and reproducible prioritization. Its modular design enables the incorporation of newly harmonized data types and iterative refinement as datasets evolve, allowing re-scoring under alternative therapeutic or safety assumptions, ensuring continuous methodological adaptation.

Considered improvements to our target identification framework include cross-technology meta-analyses and cross-validation to strengthen transcriptomic candidate identification. Additionally, the integration of protein-level expression data, such as mass spectrometry data, should be prioritized to complement gene-level expression data. This added layer of information would enhance the accuracy of biological signals and improve translational relevance, given that mRNA abundance serves only as an indirect proxy for protein expression.

Additionally, the modular architecture enables deployment beyond CRC. To date, the pipeline has been applied across 18 indications, supporting both indication-wide and subtype-informed analyses. Once datasets are harmonized, scoring and calibration are automated, accelerating hypothesis generation timelines. However, contextual interpretation of prioritized targets, including detailed literature review and comparison with discontinued or established programs, remains partially manual and represents an area for further automation through an agentic AI approach, for example.

In conclusion, this framework establishes an integrative data-driven strategy for systematic therapeutic target prioritization in oncology. By leveraging large-scale in-silico analyses, it enables scalable and flexible exploration of candidate targets, helping to de-risk downstream experimental development and supporting iterative lab-in-the-loop refinement.

## SUPPLEMENTARY MATERIAL

### Supplementary tables

**Table S1:** List and links to microarray datasets.

|  |
| --- |
| GSE100179 |
| GSE20916 |
| GSE23194 |
| GSE37283 |
| GSE37364 |
| GSE4183 |
| GSE44076 |
| GSE62932 |
| GSE100480 |
| GSE101472 |
| GSE101896 |
| GSE103340 |
| GSE106584 |
| GSE109057 |
| GSE110224 |
| GSE113513 |
| GSE119409 |
| GSE122183 |
| GSE123390 |
| GSE13067 |
| GSE13294 |
| GSE14333 |
| GSE143985 |
| GSE15960 |
| GSE161158 |
| GSE170999 |
| GSE178120 |
| GSE18088 |
| GSE18105 |
| GSE18549 |
| GSE19862 |
| GSE21510 |
| GSE23878 |
| GSE26027 |
| GSE26906 |
| GSE27157 |
| GSE27544 |
| GSE27854 |
| GSE28702 |
| GSE29621 |
| GSE31595 |
| GSE32323 |
| GSE33113 |
| GSE34489 |
| GSE35144 |
| GSE35452 |
| GSE35896 |
| GSE37892 |
| GSE39084 |
| GSE39582 |
| GSE39923 |
| GSE40367 |
| GSE41328 |
| GSE41568 |
| GSE46824 |
| GSE46862 |
| GSE52735 |
| GSE54483 |
| GSE60331 |
| GSE60697 |
| GSE62080 |
| GSE64256 |
| GSE64857 |
| GSE65480 |
| GSE71222 |
| GSE72718 |
| GSE72968 |
| GSE72969 |
| GSE75316 |
| GSE76855 |
| GSE79959 |
| GSE81558 |
| GSE84984 |
| GSE85043 |
| GSE92921 |
| GSE93375 |
| GSE96528 |

**Table S2:**
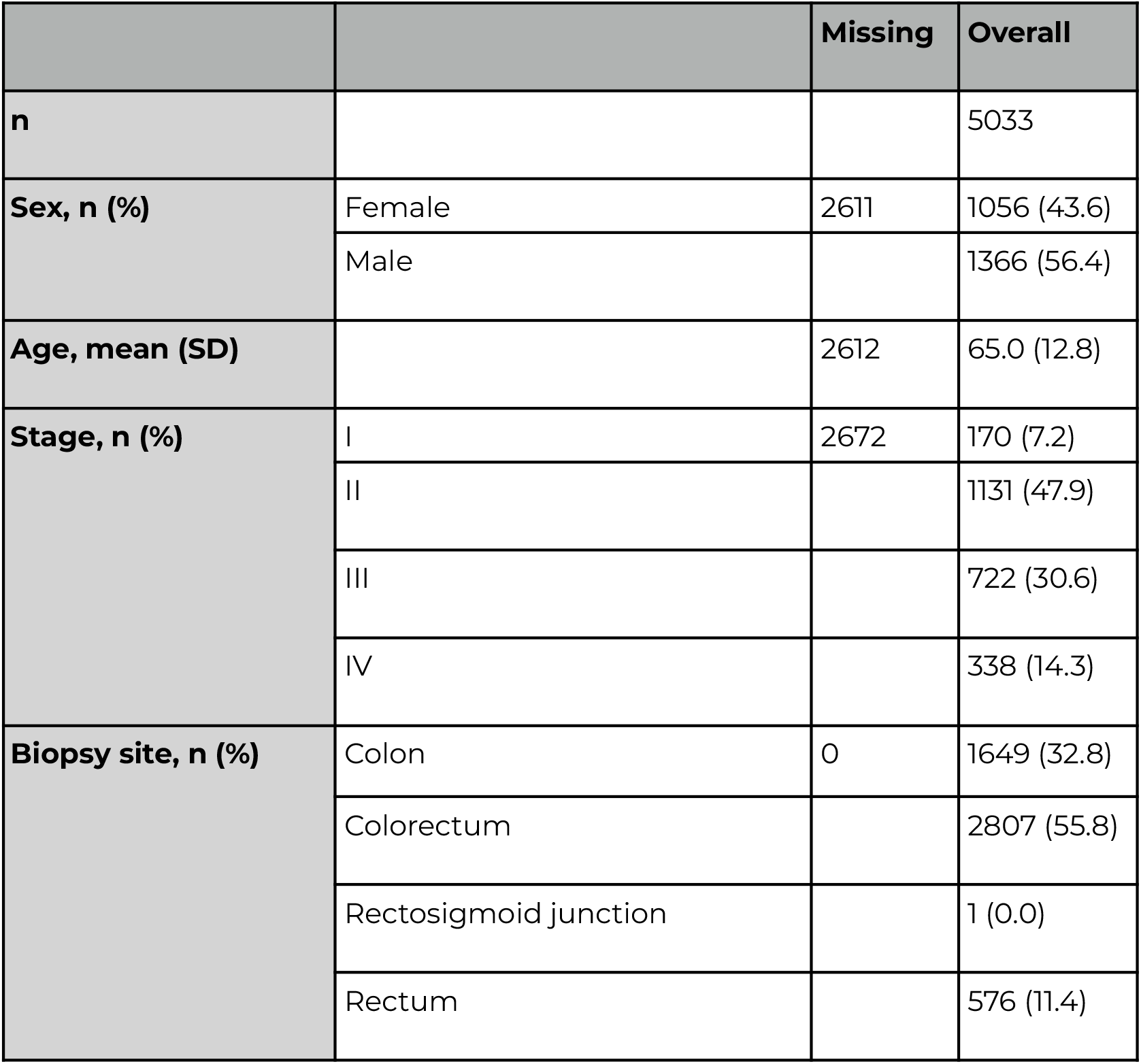
Clinical table Microarray atlas.

|  |  | Missing | Overall |
| --- | --- | --- | --- |
| **n** |  |  | 5033 |
| **Sex, n (%)** | Female | 2611 | 1056 (43.6) |
|  | Male |  | 1366 (56.4) |
| **Age, mean (SD)** |  | 2612 | 65.0 (12.8) |
| **Stage, n (%)** | I | 2672 | 170 (7.2) |
|  | II |  | 1131 (47.9) |
|  | III |  | 722 (30.6) |
|  | IV |  | 338 (14.3) |
| **Biopsy site, n (%)** | Colon | 0 | 1649 (32.8) |
|  | Colorectum |  | 2807 (55.8) |
|  | Rectosigmoid junction |  | 1 (0.0) |
|  | Rectum |  | 576 (11.4) |

**Table S3:** Clinical table TCGA COAD/READ atlas.

|  |  | Missing | Overall |
| --- | --- | --- | --- |
| **n** |  |  | 637 |
| **Age, mean (SD)** |  | 3 | 66.4 (12.7) |
| **Sex, n (%)** | Female | 6 | 294 (46.6) |
|  | Male |  | 337 (53.4) |
| **Stage, n (%)** | I | 24 | 108 (17.6) |
| **Stage, n (%)** | II |  | 233 (38.0) |
|  | III |  | 183 (29.9) |
|  | IV |  | 89 (14.5) |
| **Biopsy site, n (%)** | Colon | 4 | 467 (73.8) |
|  | Connective, subcutaneous and other soft tissues |  | 2 (0.3) |
|  | Rectosigmoid junction |  | 76 (12.0) |
|  | Rectum |  | 88 (13.9) |

**Table S4:** Target-indication pairs used for the FDA-approved benchmark.

| Target Name | Indication |
| --- | --- |
| CD19 | ALL |
| CD22 | ALL |
| CD33 | AML |
| ERBB2 | Breast |
| TACSTD2 | Breast |
| EGFR | Colon |
| EGFR | Head and neck |
| CD274 | Liver |
| CD274 | LUAD |
| EGFR | LUAD |
| ERBB2 | LUAD |
| CD274 | LUSC |
| EGFR | LUSC |
| ERBB2 | LUSC |
| FOLR1 | Ovarian |
| FOLH1 | Prostate |
| CD274 | Skin |
| ERBB2 | Stomach |

**Table S5:**
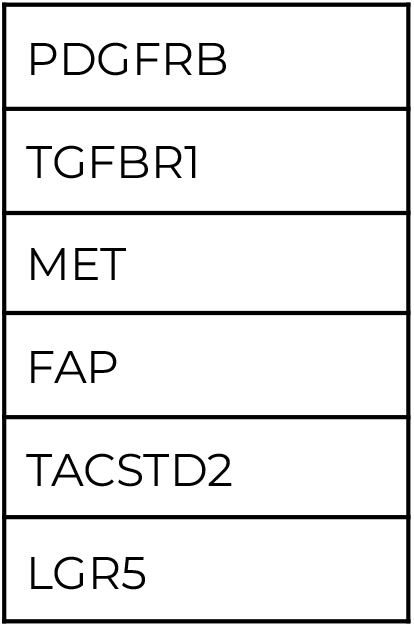
CMS4 well-established targets.

**Table S6:** List of the 11 prioritized targets.

|  |
| --- |
| FAP |
| PDPN |
| CLDN1 |
| CDH3 |
| EDNRA |
| AREG |
| THBD |
| ADAM12 |
| CDH11 |
| CD248 |
| FPR1 |

### Supplementary figures

**Figure S1:**
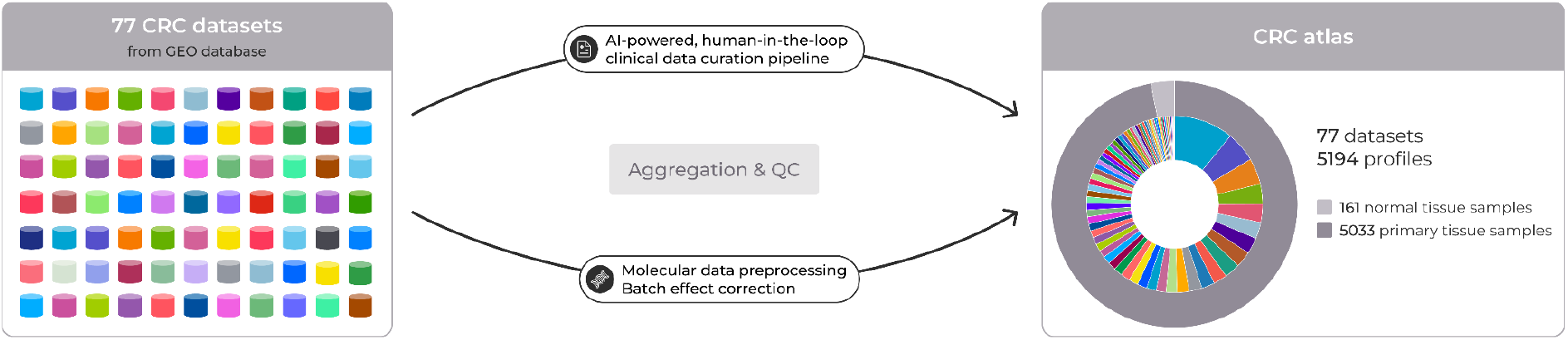
Legend: Schematic overview of the atlas construction pipeline. Individual publicly available CRC microarray datasets were retrieved, subjected to quality control, and normalized independently. Datasets were subsequently integrated and batch-corrected to generate a unified, harmonized mCRC transcriptomic atlas for downstream molecular analyses.

**Figure S2:**
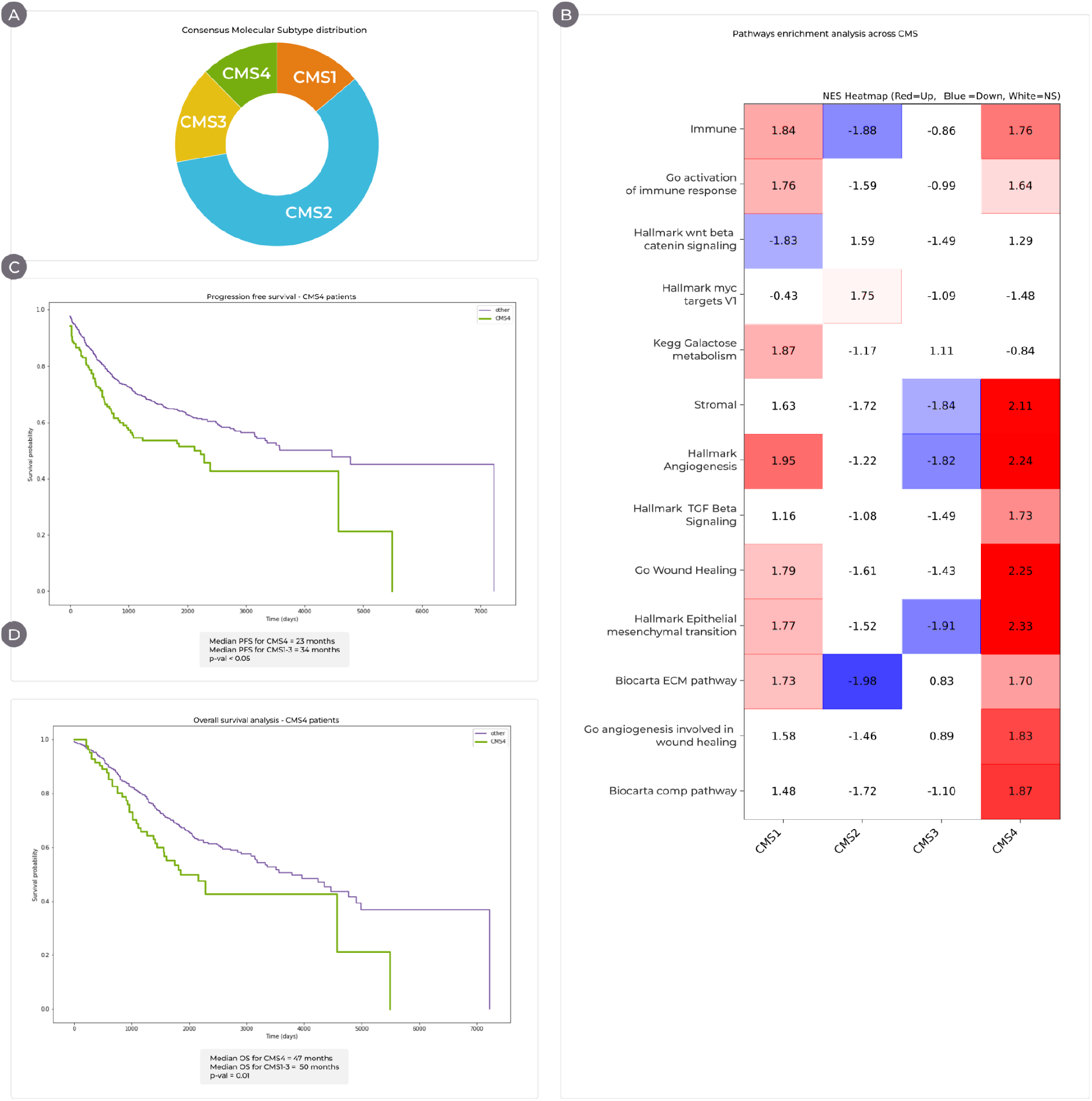
Legend: **(A)** Donut plot illustrating the distribution of consensus molecular subtypes (CMS1–4) across the metastatic colorectal cancer (mCRC) microarray atlas. **(B)** Heatmap depicting hallmark or curated pathway gene sets enriched (red) or downregulated (blue) across CMS1–4, derived from gene set enrichment analysis (GSEA) or single-sample GSEA (ssGSEA). Rows represent pathways and columns represent CMS groups. **(C–D)** Kaplan–Meier curves of progression-free survival (PFS) **(C)** and overall survival (OS) **(D)** in the CMS4 subgroup of the mCRC microarray atlas. P-value was determined by log-rank test.

